## Supplementary Information for "Weight illusions explained by efficient coding based on correlated natural statistics"

### Supplementary Text for "Weight illusions explained by efficient coding based on correlated natural statistics"

#### Variant encoding models

In the univariate case considered by Wei & Stocker[1,2], the efficient coding goal of maximizing mutual information is attained by setting Fisher Information, $J(\theta)$, to a function of the prior,

$$J(\theta) \propto p(\theta)^2. \tag{1}$$

The methods in the main text extend the efficient coding model to the case of correlated variables (mass and volume) by considering the conditional probability of mass with respect to volume, implicitly assuming that sensory evidence for volume contributes to the efficient representation of mass but is encoded independently. Here we consider how the efficient coding principle could be extended to joint encoding of the two variables with respect to their prior distribution.

##### Factorized density model

The equivalent constraint to Eq. 1 for the multivariate case is

$$\det \mathbf{J}(\boldsymbol{\theta}) \propto p(\boldsymbol{\theta})^2, \tag{2}$$

but note this does not fully constrain the Fisher Information matrix, with the implication that there is more than one encoding of a combination of variables that satisfies the efficient coding constraint. Here we present one formulation of the model that satisfies this equation, based on factorizing the prior distribution into independent components (see Barlow & Foldiak[3], for theoretical background).

We make a change of basis,

$$\mathbf{x} = \begin{bmatrix} x \\ y \end{bmatrix} = \mathbf{S}^{-1}\mathbf{R}^{-1}\left(\begin{bmatrix} m \\ v \end{bmatrix} - \begin{bmatrix} \mu_m \\ \mu_v \end{bmatrix}\right), \tag{3}$$

based on the eigendecomposition of the prior's covariance matrix,

$$\boldsymbol{\Sigma} = \mathbf{RSSR}^{-1}, \tag{4}$$

such that prior probabilities $p(x) = \phi(x)$ and $p(y) = \phi(y)$ are independent standard normal distributions, with $p(x, y) = p(x)p(y)$, and prior densities in the two bases related through

$$p_{M,V}(m, v) = p_{X,Y}(x, y) \det \mathbf{D}, \tag{5}$$

where $\mathbf{D} = \frac{\partial(x,y)}{\partial(m,v)}$ is the Jacobian of the transformation which in this case is simply $\mathbf{D} = \mathbf{RS}$. Setting Fisher Information for each independent variable according to the univariate constraint (Eq. 1) leads to the joint Fisher Information matrix,

$$\mathbf{J}_{XY} \propto \begin{bmatrix} p(x)^2 & 0 \\ 0 & p(y)^2 \end{bmatrix} \tag{6}$$

$$\implies \det \mathbf{J}_{XY} \propto p(x)^2 p(y)^2 = p(x, y)^2. \tag{7}$$

The FI matrix for the original variables is

$$\mathbf{J}_{MV} = \mathbf{D}^{\mathrm{T}} \mathbf{J}_{XY} \mathbf{D} \tag{8}$$

$$\implies \det \mathbf{J}_{MV} = (\det \mathbf{D})^2 \det \mathbf{J}_{XY} \tag{9}$$

$$\propto (\det \mathbf{D})^2 p(x,y)^2 \tag{10}$$

$$\implies \det \mathbf{J}_{MV} \propto p(m,v)^2, \tag{11}$$

satisfying Eq. 2.

Prior probabilities of $x$ and $y$ are mutually independent, so we can base predictions for efficient encoding of $x$ (or equivalently $y$) on the univariate case, i.e.,

$$b_x(x) \propto \frac{d}{dx}\frac{1}{p(x)^2} = -2\frac{p'(x)}{p(x)^3} = 2\frac{x}{\phi(x)^2} \tag{12}$$

$$\implies b'_x(x) \propto \frac{2+4x^2}{\phi(x)^2}, \tag{13}$$

and s.d.

$$\hat{\sigma}_x(x) \propto \frac{1+b'_x(x)}{p(x)} = \frac{1}{\phi(x)} + \frac{2+4x^2}{\phi(x)^3}. \tag{14}$$

Bias in the original variables can be obtained as

$$\begin{bmatrix} b_m(m) \\ b_v(v) \end{bmatrix} = \mathbf{RS} \begin{bmatrix} b_x(x) \\ b_y(y) \end{bmatrix}, \tag{15}$$

and s.d. from

$$\mathrm{Cov}(m,v) = \begin{bmatrix} \hat{\sigma}^2_m(m) & \hat{\sigma}(m,v) \\ \hat{\sigma}(m,v) & \hat{\sigma}^2(v) \end{bmatrix} = \mathbf{RS} \begin{bmatrix} \hat{\sigma}^2_x(x) & 0 \\ 0 & \hat{\sigma}^2_y(y) \end{bmatrix} \mathbf{SR}^{\mathrm{T}}. \tag{16}$$

Predictions for this model are illustrated in Fig. S2B. Predictions for the main (conditional density) model are duplicated in Fig. S2A for comparison.

##### Joint density model

The model in the main text allocates resources for encoding object mass according to the conditional prior probability of mass given object volume, i.e. $J_M \propto p(m|v)$. This is an appropriate solution if the total resources available to encode mass are constant irrespective of volume, e.g., the case of distributing preferred values in a neural population code. This also allows for the simple coding scheme discussed in the main text, in which a population with fixed tuning encodes deviations from the expected mass given object volume.

An alternative is that the total resource for encoding mass can update according to volume, for example, by gain modulation in a population code where the goal is to maximize mutual information under an energetic constraint. In such cases, an appropriate solution would be to allocate resources according to the joint prior probability of mass and volume, $J_M \propto p(m,v)$, thereby conserving activity for those objects with the most frequently encountered combinations of properties. Predictions for this model are illustrated in Fig. S2C.

##### Model comparison

Predictions based on maximum likelihood parameters of the factorized (green) and joint density (blue) models for biases and SD are shown in Fig. S3A & B. Fits were broadly similar to the conditional density model used in the main text and captured all the same qualitative features of the data. Formal model comparison using AIC indicated that the factorized density model (green in Fig. S3C) in most cases fit less well than the joint density model (blue), which performed very similarly to the conditional density model (baseline, red). The best fitting model was the conditional density model for 53% of participants (16/30), the joint density model for 37% (11/30) and the factorized density model for 10% (3/30).

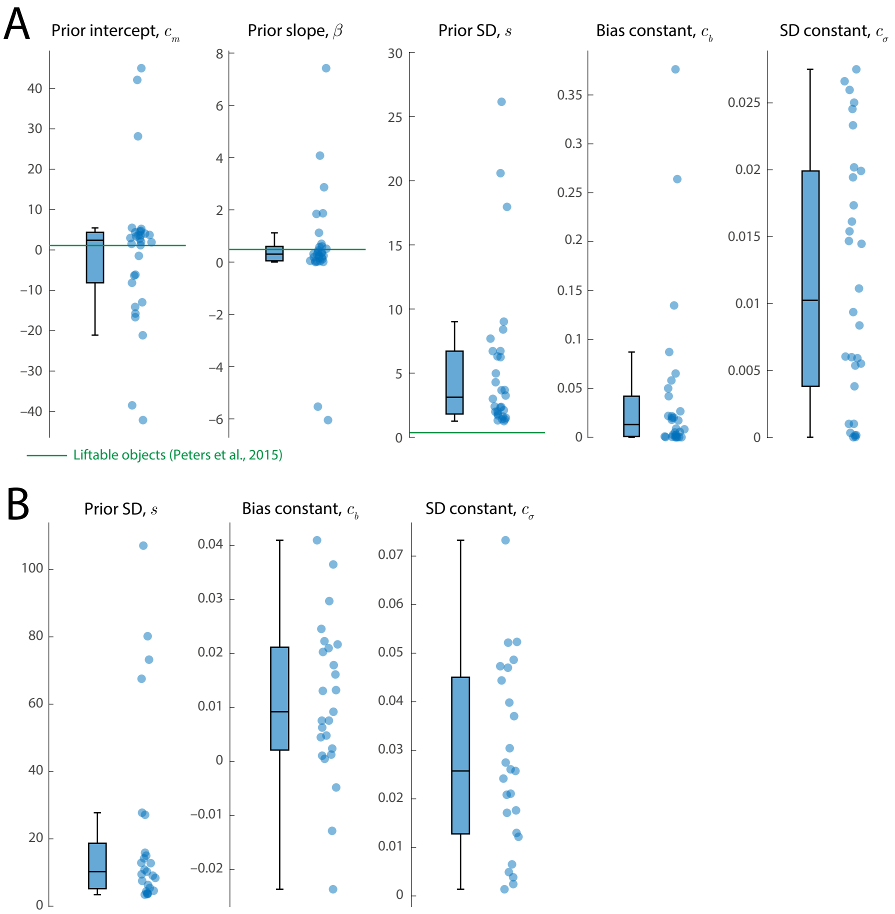


Supplementary Figure 1: Model parameters. (A) Maximum likelihood parameters for the SWI model. Datapoints correspond to individual participants. For comparison, green lines indicate values of the first three model parameters that would specify a prior distribution that matched the distribution of liftable objects sampled by Peters *et al.*[4] and illustrated in Fig. 1A. (B) Maximum likelihood parameters for the MWI model. Box plots: centre-line, median; box limits, upper and lower quartiles; whiskers, 1.5× interquartile range.

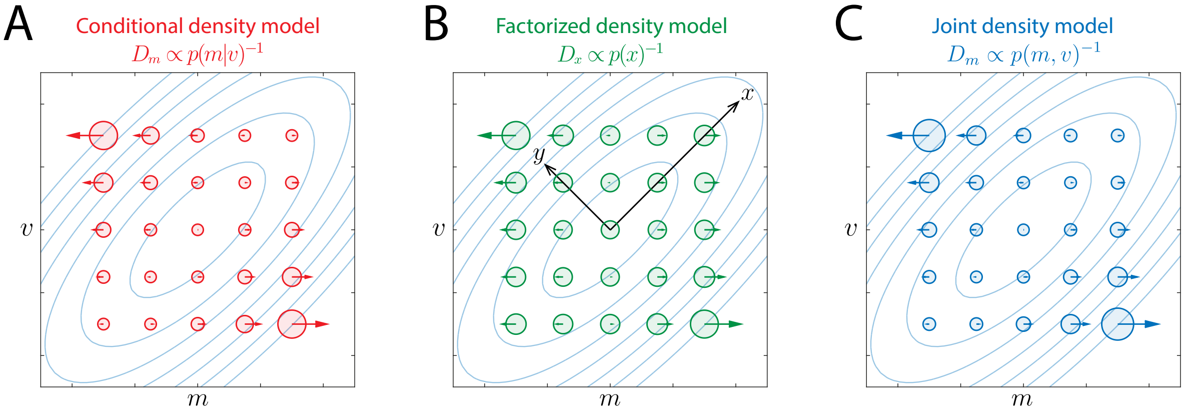


Supplementary Figure 2: Predictions of variant encoding models. Predicted SD (circle diameters) and bias (arrows) in estimates of log-mass for objects with a range of true log-volumes ($v$) and log-masses ($m$), for (A) the model in the main text, where object mass is efficiently encoded with respect to its conditional prior probability given object volume; (B) a model where object mass and volume are jointly efficiently encoded with respect to the eigenvectors ($x$ and $y$) of the prior distribution; (C) a model where object mass is efficiently encoded with respect to the joint prior probability of mass and volume.

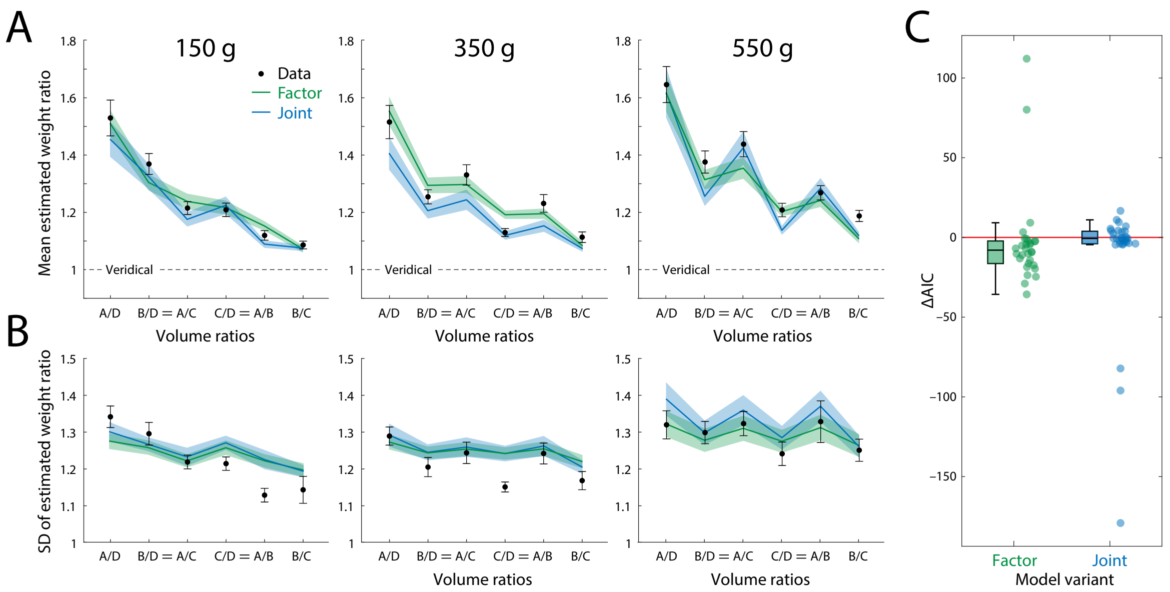


Supplementary Figure 3: Fits of variant encoding models. (A) Mean weight ratio reported by participants (black) for each pairing of lifted objects, with predictions of factorized (green) and joint density (blue) models. Different panels show results for three sets of objects with different common masses. Error bars and shading indicate ±1 SEM. (B) Within-participant SD of weight ratios, plotted as in A. (C) Model comparison results plotted as AIC values for factorized (green) and joint density (blue) models relative to conditional density model. Positive values (above baseline) indicate a better fit for the variant model. Data points correspond to individual participants.

#### References


1. Wei, X.-X. & Stocker, A. A. A Bayesian Observer Model Constrained by Efficient Coding Can Explain 'anti-Bayesian' Percepts. *Nature Neuroscience* **18,** 1509–1517. ISSN: 1546-1726 (2015).
2. Wei, X.-X. & Stocker, A. A. Lawful Relation between Perceptual Bias and Discriminability. *Proceedings of the National Academy of Sciences,* 201619153 (2017).
3. Barlow, H. B. & Foldiak, P. in *Miall C, Durbin RM, Mitchison GJ, Eds. The Computing Neuron* 54–72 (Wokingham, United Kingdom: Addison-Wesley, 1989).
4. Peters, M. A. K., Balzer, J. & Shams, L. Smaller = Denser, and the Brain Knows It: Natural Statistics of Object Density Shape Weight Expectations. *PLOS ONE* **10,** e0119794. ISSN: 1932-6203 (2015).
